# A Synergistic Anti-Cancer FAK and HDAC Inhibitor Combination Discovered by a Novel Chemical-Genetic High-Content Phenotypic Screen

**DOI:** 10.1101/590802

**Authors:** John C. Dawson, Bryan Serrels, Adam Byron, Morwenna Muir, Ashraff Makda, Amaya García-Muñoz, Alex von Kriegsheim, Neil O. Carragher, Margaret C. Frame

## Abstract

We mutated the Focal Adhesion Kinase (FAK) catalytic domain to inhibit binding of the chaperone, Cdc37 and ATP, mimicking the actions of a FAK kinase inhibitor. We re-expressed mutant and wild-type FAK in squamous cell carcinoma (SCC) cells from which endogenous FAK had been deleted, genetically fixing one axis of a FAK inhibitor combination high-content phenotypic screen to discover drugs that may synergize with FAK inhibitors. HDAC inhibitors represented the major class of compounds that potently induced multi-parametric phenotypic changes when FAK was rendered kinase-defective, or inhibited pharmacologically in SCC cells. Indeed, combined FAK and HDAC inhibitors arrest proliferation and induce apoptosis in a sub-set of cancer cell lines in vitro and efficiently inhibits tumor growth in vivo. Mechanistically, HDAC inhibitors potentiate inhibitor-induced FAK inactivation and reverses FAK-associated nuclear YAP in sensitive cancer cell lines. Here we report the discovery of a new clinically actionable synergistic combination between FAK and HDAC inhibitors.

**Significance:** We describe a chemical-genetic, high-content phenotypic screening approach to discover effective anti-cancer combinations of targeted therapeutics through which we identified a novel, synergistic and clinically actionable, combination of FAK and HDAC inhibitors.

## Introduction

Focal adhesion kinase (FAK) is a non-receptor tyrosine kinase that is frequently over-expressed in cancer, where it functions downstream of integrins and growth factor receptors to regulate a diverse range of cellular functions, including survival, proliferation, adhesion, migration, angiogenesis, stemness, and more recently, cytokine expression (1,2). FAK plays a pivotal role in transducing signals from the plasma membrane and the nucleus (2–8). It is recognized as a high-value druggable target for cancer therapy, resulting in the development of multiple FAK kinase inhibitors that are being tested in clinical trials (reviewed in Lee et al., (1)). We have argued that FAK inhibitors may best be used in combination with other agents because FAK is part of key survival pathways, including when cancer cells are exposed to stress, including when treated with anti-cancer agents (4,5,9). Here we report a new chemical-genetic phenotypic screening approach to identify combinations between FAK, and other targeted, inhibitors.

The multiparametric, high-content phenotypic screening approach we took enables unbiased “morphometric profiling” of compound activity and quantification of functional similarities and dissimilarities between drug mechanisms of action (MOAs) across distinct cell types (10–13). Integration of information-rich phenotypic assays (as opposed to information-poor “black-box” phenotypic assays) with the latest advances in multiparametric image analysis, multivariate statistics, and new image informatics tools, permits the classification of compound-induced morphometric phenotypes, comparing cancer cells that differed only in FAK activity status. Phenotypic “fingerprints” derived from multiparametric, high-content morphometric measurements were used to identify clusters enriched with specific compound MOA and target classes which, when perturbed, induced a characteristic phenotypic response between cell lines that differed only in FAK activity.

In this way, we combined high-content phenotypic screening with cancer cells genetically-modified to express a new discriminatory FAK mutant protein (described below) to identify synergistic FAK inhibitor combinations. We generated a novel kinase-defective FAK protein (FAK-G431A,F433A) that abolished binding of the Cdc37 chaperone protein, and likely ATP. This FAK-G431A,F433A mutant behaved, in many respects, including Cdc37 binding, more like FAK in the presence of a small-molecule FAK inhibitor when it was compared to a conventional kinase-defective FAK mutant (FAK-KD). We re-expressed FAK-G431A,F433A in squamous cell carcinoma (SCC) cells from which the FAK gene (*Ptk2*) had been previously deleted by Cre-lox-mediated recombination (as we have described before (2,5,14)), allowing us to compare the loss of FAK’s kinase activity with cells that do not express FAK (FAK−/−) or that re-express wild-type FAK (FAK-WT). This allowed us to genetically fix one axis of a multiparametric phenotypic screen; from this, we identified distinct phenotypic responses between cells expressing FAK-WT and FAK-G431A,F433A after treatment with histone deacetylase (HDAC) inhibitors, indicative of synergism. This synergy was confirmed by combining pharmacological FAK kinase and HDAC inhibitors in subsequent dose-ratio combination matrix experiments. In screening multiple cell lines, we identified a sub-set of cancer cells in which FAK and HDAC inhibitors synergized to inhibit proliferation. In these, HDAC inhibitors enhanced FAK inhibitor-induced FAK inactivation as judged by FAK-Y397 autophosphorylation, and this was linked to enhanced nuclear exclusion of YAP in vitro and in vivo, suggesting that combining FAK and HDAC inhibitors promotes dysregulation of Hippo/YAP signaling.

## Materials and Methods

VS-4718 (#S7653), vorinostat (#S1047), panobinostat (#S1030), staurosporine (#S1421) and paclitaxel (#S1150) were purchased from Selleck Chemicals. Compound libraries were from BioAscent. Antibodies used were purchased from Cell Signaling Technologies (FAK pY397, #3283; FAK, #3285; GAPDH, #2118; YAP, #14074; YAP pS127, #13008; Histone H3, #9715; Histone H3 K-Ac Lys 56, #4243; Cdc37, #4793; Cdc37 pS13, #13248; Src, #2109) or AbCam (YAP, #EP1674Y).

### Cell Lines

The SCC FAK cell model was generated and isolated and grown as previously described (6). SCC FAK−/− cells stably re-expressing FAK-WT, FAK-KD, or FAK-G431A,F433A were maintained under selection using 0.25 mg/mL hygromycin (Invivogen, #ant-hg-1). Human cell lines were cultured in RPMI (Sigma-Aldrich, #R0883) supplemented with 2 mmol/L L-glutamine (ThermoFisher Scientific, #25030081) and 10% FBS. Human cell lines were purchased from ATCC and esophageal cells were a gift of T. Hupp (University of Edinburgh). Cell lines were routinely tested for mycoplasma (twice yearly) and cell line authenticity confirmed by STR profiling where appropriate.

### Subcutaneous Tumor Growth

Experiments involving animals were carried out in accordance with the UK Coordinating Committee on Cancer Research guidelines by approved protocol (HO PL 70/8897). SCC FAK-WT cells (0.25 × 10^6^) and A549 cells (5 × 10^6^) were injected subcutaneously on each flank of CD1-nude mice (5 per group). Established tumor-bearing mice were randomized into treatment groups and dosed with panobinostat (5 mg/kg) 5 days out of 7 by IP injection (for a total of 14 injections) and twice daily with VS-4718 by oral gavage. Panobinostat was dissolved in DMSO and diluted in 10% PEG400/1% Tween-80 immediately prior to injection. VS-4718 was prepared in 0.5% hydroxypropyl methylcellulose. Tumor growth was monitored by caliper measurements and tumor volume calculated using the formula *V* = (*W*^2^ × *L*)/2, where *V* is tumor volume, *W* is tumor width, and *L* is tumor length. Animals were sacrificed when tumors reached their maximum allowable size or when tumor ulceration occurred.

### Immunoprecipitation and Immunoblot Analysis

Cell lysates were prepared for immunoblotting and immunoprecipitation as previously described (3,7).

### Reverse Phase Protein Array (RPPA) Analysis

Flo1 cell lysates from 24 hour drug treated cells were isolated in RIPA as described above and processed for RPPA (35) by the IGMM Protein and Antibody Microarray Facility (Cancer Research UK Edinburgh Centre, University of Edinburgh).

### Immunofluorescence

Cells were grown on glass coverslips and processed as previously described (5).

### High-Content Imaging

#### FAK Mutant Combination Screen

SCC cells were seeded in collagen I-coated 384-well optical bottom plates (Greiner; #781091) at 300 cells per well. Cells were cultured for 24 hours before addition of compounds using a BioMek FX liquid handling station for a further 24 hours. Plates were fixed with 4% paraformaldehyde in PBS for 20 minutes followed by washing with PBS twice and incubated with blocking solution (0.2% Triton X-100, 1% BSA in PBS) for 30 minutes. Plates were incubated with Alexa Fluor 488-labeled phalloidin (1:250; Cell Signaling Technology; #8878) and Hoechst 33345 (2 μg/mL; Molecular Probes; #H1399) made up in blocking solution for 50 minutes. Finally, an equal volume of HCS Cell Mask deep red (1:75,000; Molecular Probes; #H32721) diluted in blocking solution was added for 10 minutes. Plates were washed with PBS and plates sealed. Six images per well were acquired using an ImageXpress Micro XLS widefield microscope (Molecular Devices) with a 20x S Plan Fluor objective, with typically 150–200 cells per image of control wells.

Each plate contained 32 wells treated with 0.1% DMSO as a negative control and 16 wells treated with paclitaxel (300 nmol/L) and 16 wells with staurosporine (300 nmol/L) as positive controls.

#### YAP Localization

Cells in 384-well plates were fixed with 4% paraformaldehyde for 20 minutes. Plates were washed with PBS and cells permeabilized and blocked (0.3% Triton X-100, 2.5% FBS in PBS) for 30 minutes. Plates were immunolabeled with anti-YAP antibody (AbCam, #EP1674Y) diluted in blocking buffer and incubated at 4°C overnight. Plates were washed with PBS and labeled with anti-rabbit Alexa Fluor 594-conjugated secondary antibody and Hoechst (2 μg/mL) diluted in blocking buffer. Plates were washed in PBS and sealed. Nine images per well were acquired using an ImageXpress Micro XLS widefield microscope with a 20x S Plan Fluor objective, with typically 100–200 cells per image and 6 replicate wells per condition per plate.

#### Image Analysis

Images were quantified using CellProfiler (36) or MetaXpress software (Molecular Devices). For CellProfiler, briefly, for all image analysis pipelines, nuclei were identified using the ‘Identify Primary Objects’ module using an Otsu threshold followed by one of the following methods. For the FAK mutant combination screen, the phalloidin and HCS Cell Mask channels were combined together and used to derive a ‘cell’ object using the ‘propagate’ function in the ‘Identify Secondary Objects’ module. Finally, a mask was created using an Otsu threshold on the HCS Cell Mask channel, which was subtracted from the ‘cell’ mask to give cellular ‘protrusions’. For YAP localization, YAP immunostaining was used to derive a ‘cell’ object by expanding the nuclear object by 50 pixels without joining any other object in the ‘Identify Secondary Objects’ module. The ‘nuclear’ object was subtracted from the ‘cell’ object to give the ‘cytoplasm’. The fluorescent intensity of anti-YAP immunocytochemistry in the nuclear and cytoplasmic regions was measured per cell. For cell cycle analysis, images of Hoechst-labeled nuclei were analyzed using the ‘Cell Cycle’ module in the MetaXpress software.

#### Data Analysis

Data were analyzed using the TIBCO Spotfire High Content Profiler application (PerkinElmer). For the FAK mutant combination screen, cellular measurements were aggregated to image median values and then to well level. Plates were median normalized to the DMSO controls of each individual plate and then each plate was normalized to the global median of the DMSO wells and Z-scores were calculated. Wells that contained fewer than 50 cells were excluded. To identify differential effects between FAK-WT and FAK-mutant (FAK-G431A,F433A or FAK−/−) cells, the FAK-WT value for each treatment was subtracted from each FAK mutant value. Data was then analyzed using PCA, and a Mahalanobis distance was used to identify differentially active compounds. Hit compounds were selected as having a Mahalanobis distance Z-score of greater than 2.

### Live-Cell Kinetic Analysis of Cell Cycle

SCC FAK-WT cells stably expressing the FUCCI cell cycle reporter (3) were seeded on 96-well collagen I-coated plates. Cells were treated with drug combinations in a dose matrix and plates monitored in an Incucyte Zoom microscope (Essen BioScience) at 37°C over 5 days. Images were analyzed using the inbuilt analysis software.

### Apoptosis Assay

Cells were seeded in 96-well plates with IncuCyte^®^ Caspase-3/7 green apoptosis assay reagent reagent (Essen BioScience; #4440) and treated with compounds. Plates were imaged in an Incucyte Zoom, acquiring images every 3 hours over a 72 hour period using the ‘phase’ and ‘green’ channels. Images were analyzed using the Incucyte Zoom software, and apoptosis is expressed as a percentage of the population.

### Generation of Spheroids

Cells were seeded in 96-well ultra-low attachment “U” bottom plates (Corning; #7007) at 2000 cells per well in normal growth medium and centrifuged at 1000 × *g*. Cells were allowed to form a single spheroid per well over 3 days, after which they were drug treated and monitored over 14 days for using an ImageXpress Micro XLS widefield microscope. For assessment of spheroid viability, spheroids were incubated with calcein AM (4 μmol/L final concentration; Molecular Probes; #C3099) for 1 hour prior to imaging.

### Immunohistochemistry

For immunohistochemistry, reagents were from DAKO using standard procedures. Briefly, tumor sections (4 μm thick) were rehydrated, and antigen retrieval was performed using boiling citrate buffer (pH 6.0) for 5 minutes. Endogenous peroxidase activity was blocked using peroxidase blocking solution (#S2023) and then sections were incubated with serum-free blocking solution (#X0909). Sections were incubated with diluted primary antibody overnight at 4°C. Sections were washed in TBS and incubated with secondary (#P0448) for 30 minutes. Sections were washed in TBS and incubated with DAB reagent (#K3468) for 5 minutes, and finally, sections were counter-stained with Eosin, dehydrated, and mounted using DPX mounting medium (#44581).

### Cell Viability Assay

Cells were seeded in 96-well plates at 2000 cells per well and incubated for 24 hours before treatment. Cells were incubated with compounds for 72 hours, untreated cells were incubated with 0.1% DMSO. Alamar Blue (Invitrogen; #DAL1025) was added to each well and plates incubated for 3 hours. Fluorescence emission was read on an EnVision 2101 multilabel plate reader (PerkinElmer; excitation = 540 nm, emission = 590 nm). All conditions were normalized to plate DMSO control wells.

### Colony Formation Assay

Single cells (1000 per well) were seeded in six-well plates, treated with drugs, and allowed to grow over 7–10 days. Plates were washed with PBS and colonies stained with Coomassie Blue (0.25 g/L Coomassie Brilliant Blue R-250 (#27816) made up in 45% water, 45% acetic acid, and 10% methanol). The colony area covering the well was quantified in ImageJ.

### Synergy Calculation

Normalized measurements were averaged (*n* from ≥ 3 independent experiments) and analyzed using SynergyFinder (37) using the Bliss, Loewe, and ZIP synergy models.

### Metabolic Labeling

SCC cells were cultured in SILAC MEM (DC Biosciences) containing either L-[U-^13^C_6_]arginine and L-[^2^H_4_]lysine (R6K4; medium label) or unlabeled L-arginine and unlabeled L-lysine (R0K0; light label) for more than six cell doublings. At time ***t*** = 0 h, cells were washed twice in warm PBS, and either 1 × 10^6^ light-labeled cells were re-plated in light-labeled medium or 1 × 10^6^ medium-labeled cells were replated in SILAC MEM containing L-[U-^13^C_6_,^15^N_4_]arginine and L-[U-^13^C_6_,^15^N_2_]lysine (R10K8; heavy label). SILAC MEM was supplemented with 10% dialyzed FBS, 2 mmol/L L-glutamine, 1 mmol/L sodium pyruvate, MEM vitamins, and MEM non-essential amino acids throughout the experiments. Cells were harvested and lysed in RIPA buffer (as above) at *t* = 0.5, 6, 20, 45, and 70 h. Light-labeled cell lysates were mixed with medium-/heavy-labeled cell lysates at 1:1 ratio by total protein amount, mixed lysates were resolved by polyacrylamide gel electrophoresis, and gels were stained with InstantBlue (Expedeon).

### Mass Spectrometry

#### Analysis of Immune Complexes

For analysis of immune complexes using label-free mass spectrometry, captured proteins were subjected to on-bead proteolytic digestion, desalting, and liquid chromatography–tandem mass spectrometry, as previously described (38). Peptide and protein false discovery rates were set to 1%. Mean label-free intensities were calculated from technical duplicate mass spectrometry runs for each of three biological replicates per experimental condition.

#### FAK Turnover Profiling

For FAK turnover profiling using SILAC-based mass spectrometry, gel portions spanning the molecular weight range 100–150 kDa of resolved mixed lysates (see above) were excised and subjected to in-gel proteolytic digestion, as previously described (39). Peptides were analyzed by liquid chromatography–tandem mass spectrometry, as previously described (40). Peptide and protein false discovery rates were set to 1%. SILAC ratios (medium label/light label, heavy label/light label, and heavy label/medium label) were calculated for each of three biological replicates per experimental condition. FAK synthesis and degradation curves were determined from normalized SILAC ratio profiles (heavy label/light label and medium label/light label, respectively) by nonlinear regression and plotted as means ± SEM with best fit curves and 95% confidence interval (CI) bands using Prism (GraphPad). We modeled FAK synthesis as a one-phase association curve for which *y* = *y*_0_ + (*y*_∞_ − *y*_0_) × (1 − e^−*kt*^), where *y*_0_ is the normalized SILAC ratio (heavy label/light label) at time *t* = 0, *y*_∞_ is the normalized SILAC ratio (heavy label/light label) at *t* = ∞, and *k* is the rate constant. We modeled FAK degradation as a one-phase decay curve for which *y* = (*y*_0_ − *y*_∞_) × e^−*kt*^ + *y*_∞_, where *y*_0_ is the normalized SILAC ratio (medium label/light label) at *t* = 0, *y*_∞_ is the normalized SILAC ratio (medium label/light label) at *t* = ∞, and *k* is the rate constant. Inferred FAK turnover time was calculated as the intersection between the synthesis and degradation curves. Statistical significance of synthesis and degradation curves was determined by extra sum-of-squares *F* test (α = 0.025; degradation, *F* = 0.225, DF_n_ = 6, DF_d_ = 320; synthesis, *F* = 0.170, DF_n_ = 6, DF_d_ = 319). Statistical significance of turnover times was determined by two-tailed Mann-Whitney *U* test (α = 0.05).

## Results

### Characterization of a Novel Kinase-Defective FAK that Mimics Pharmacological Inhibition

We initially identified a novel FAK binding partner, the protein kinase chaperone Cdc37 (Fig. 1A; Supplementary Fig. S1A), and found that mutations of glycine-431 and phenylalanine-433 of FAK to alanine (FAK-G431A,F433A) abolished Cdc37 binding, while a previously described kinase-inactivating mutation of FAK (K454R, FAK-KD (15)) did not (Fig. 1A). This mimicked the effect of an ATP-competitive FAK kinase inhibitor, VS-4718, which blocked Cdc37 binding to FAK-WT (Fig. 1B). We confirmed that FAK-G431A,F433A was completely defective in kinase activity as measured by lack of any visible autophosphorylation at FAK-Y397 (Fig. 1C), presumably because it disturbs the ATP binding site. Moreover, FAK-G431A,F433A impairs binding to Src (Fig. 1D), although it does not impair binding to kinase-independent partners, including the scaffold protein RACK1 (Supplementary Fig. S1B (8)). Further characterization revealed that Cdc37 co-localized with FAK at sites of focal adhesions, and loss of Cdc37 binding did not alter FAK’s localization to focal adhesions (Fig. 1E and F). If the FAK-G431A,F433A mutant was to be efficiently used to mimic effects of pharmacological FAK kinase inhibitors, it was important to determine whether the stability of FAK was affected by loss of Cdc37 binding, since Cdc37 is an Hsp90 co-chaperone protein that may influence protein turnover (16). Therefore, we quantified protein synthesis and degradation in SCC cells expressing FAK-WT and FAK-G431A,F433A using dynamic stable isotope labeling of amino acids in cell culture (SILAC) and mass spectrometry. We analyzed normalized SILAC ratio profiles derived from metabolically labeled FAK peptides, and inferred FAK turnover times indicated that loss of Cdc37 binding did not significantly affect FAK-G431A,F433A protein turnover when compared to FAK-WT (Fig. 1G; Supplementary Fig. S1C).

**Figure 1.**
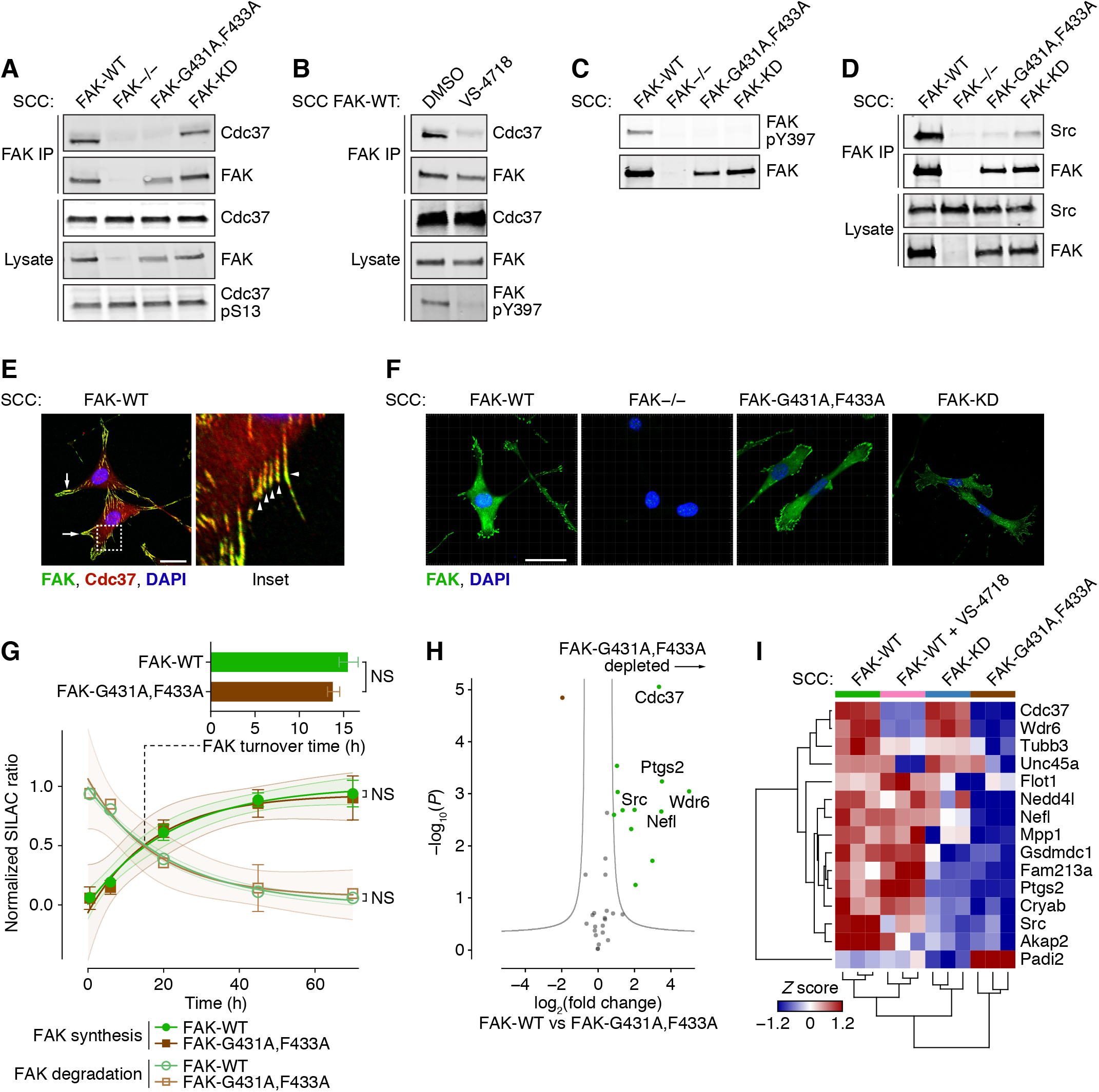
Disruption of Cdc37 chaperone binding to FAK mimics FAK kinase inhibition. **A,** IP of FAK from SCC FAK-WT, FAK−/−, FAK-G431A,F433A, and FAK-KD cell lysates, immunoblotted for FAK, Cdc37, and phosphorylated Cdc37 (S13). **B,** IP of FAK from SCC FAK-WT cells treated with DMSO (0.1%) or 250 nM VS-4718 for 24 h, immunoblotted for FAK, phosphorylated FAK (Y397), and Cdc37. **C,** Lysates from SCC FAK-WT, FAK−/−, FAK-G431A,F433A, and FAK-KD cells, immunoblotted for FAK and phosphorylated FAK (Y397). **D,** IP of FAK from SCC FAK-WT, FAK−/−, FAK-G431A,F433A, and FAK-KD cell lysates, immunoblotted for FAK and Src. **E,** SCC FAK-WT cells seeded on glass coverslips, fixed, and labeled with anti-FAK (green), anti-Cdc37 (red), and DAPI (blue). Inset (right) represents dashed boxed region of main image. Arrows and arrowheads (inset) indicate examples of regions of colocalization of FAK and Cdc37. Scale bar, 20 μm. **F,** SCC FAK-WT, FAK−/−, FAK-G431A,F433A, and FAK-KD cells seeded on glass coverslips, fixed, and labeled with anti-FAK (green) and DAPI (blue). Scale bars, 20 μm. **G,** FAK synthesis and degradation profiles quantified by metabolic labeling and mass spectrometry. FAK-WT (green) and FAK-G431A,F433A (brown) synthesis and degradation curves were determined from normalized SILAC ratio profiles by nonlinear regression and plotted as means ± SEM with best fit curves and 95% CI bands (*n* ≥ 5 peptides; representative of three independent experiments). NS, not significant (extra sum-of-squares *F* test). Bar chart inset summarizes inferred 50% protein turnover times for FAK-WT and FAK-G431A,F433A, plotted as means ± SD (*n* = 3 independent experiments). NS, not significant (two-tailed Mann–Whitney *U* test). **H,** Label-free mass spectrometric characterization of FAK-interacting proteins isolated by IP of FAK from SCC FAK-WT and FAK-G431A,F433A cell lysates. Specific FAK-binding proteins were determined versus IP from SCC FAK−/− cells (n = 3 independent experiments), satisfying *Q* < 0.05 (Student’s *t*-test with permutation-based FDR correction). Gray curves show the threshold for significant differential regulation (FDR, 5%; artificial within-groups variance, 1). Proteins satisfying *P* < 0.01 and fold change > 4 are labeled with gene names for clarity. **I,** Label-free mass spectrometric characterization of FAK-interacting proteins isolated by IP from SCC FAK-WT, FAK-KD, and FAK-G431A,F433A cell lysates and lysates from SCC FAK-WT cells treated with 250 nM VS-4718 (*n* = 3 independent experiments). Normalized label-free quantification of protein abundance for each protein was converted to a *Z* score. Differentially regulated, specific FAK-binding proteins satisfying *Q* < 0.05 (one-way ANOVA with permutation-based FDR correction) were hierarchically clustered and displayed as a heatmap. Proteins are labeled with gene names for clarity.

To assess the impact of the G431A,F433A mutations on binding partners of FAK, we isolated FAK protein complexes using immunoprecipitation and analyzed the interactomes using label-free mass spectrometry (Supplementary Fig. S1D–G). FAK-G431A,F433A exhibited impaired binding to all of the proteins for which binding to the more classical kinase-inactivating FAK-KD mutant was reduced (Fig. 1H and I; Supplementary Table S1). In addition, the FAK-G431A,F433A mutant protein interactome did not contain protein interactors that were also disrupted by the ATP-competitive FAK kinase inhibitor VS-4718, most notably Cdc37 and Wdr6 (Fig. 1H and I; Supplementary Table S1), when compared to FAK-KD, further supporting the notion that FAK-G431A,F433A both behaves like the more commonly used catalytically-inactive FAK-KD mutant and better represents the actions of pharmacologically inhibited FAK-WT protein in terms of its interactome.

### Identifying Synergistic Combinations with FAK Inhibitors

A number of FAK inhibitors are currently undergoing clinical development (1). However, it is evident from studies to date that FAK inhibitors have limited effects on cancer cell growth and survival when used as a monotherapy. In contrast, FAK’s role as a key signaling mediator of survival under conditions of cellular stress (17), including upon treatment of tumors with chemotherapy (9,18), suggests that targeting FAK in combination with other agents (that induce stress) may be beneficial. To identify potential drug combination partners for FAK inhibitors, we performed a chemical-genetic, multiparametric, high-content phenotypic screen with a small-molecule compound library and compared phenotypic responses between SCC cell lines expressing FAK-G431A,F433A, FAK-WT, or no FAK (FAK−/−). These genetically matched SCC cells differed only with regard to FAK status and the resultant consequences of modulating FAK’s kinase activity on compound response. The cells were incubated for 24 hours with an annotated compound library consisting of FDA-approved drugs and a mechanistic toolbox of kinase, epigenetic, and protease inhibitors to cover a range of biological areas (1280 and 176 compounds, respectively; Supplementary Table S2; compounds used at a final concentration of 10 μM). After compound treatment, the cells were fixed and labeled with Hoechst, phalloidin, and HCS Cell Mask to provide unbiased morphological profiling of the compound-induced phenotype. We measured multiple cellular features at the single-cell level to enable multiparametric analysis of compound-induced phenotypic responses, including principal component analysis (PCA) to interrogate major phenotypic differences between compound effects across distinct cell types. We compared the SCC cells expressing FAK-G431A,F433A with cells expressing FAK-WT and FAK-deficient (−/−) cells with FAK-WT-expressing cells (Fig. 2A).

**Figure 2.**
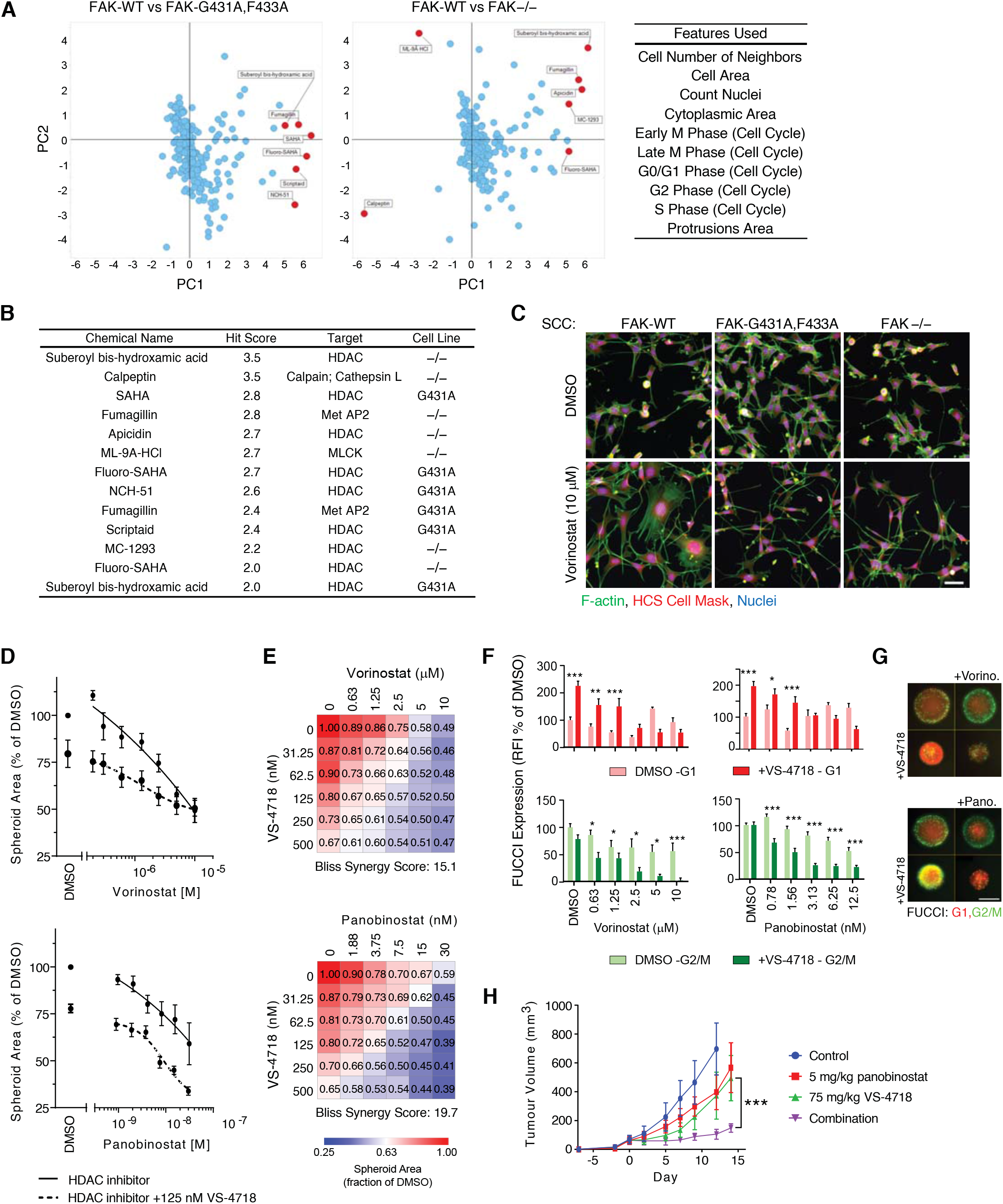
Identification of synergistic combinations with FAK kinase inhibitors. **A,** Summary of compound screen to identify drug sensitivity in the presence of reduced FAK kinase activity. PCA analysis of image-based measurements from the compound screen, centered to FAK-WT data, reveals compounds that induce a distinct phenotypic response in the SCC FAK-G431A,F433A and FAK−/− cells. Principal components (PCs) 1 and 2 account for 51.3% and 22.1% of the total variance, respectively. Cellular features used in the phenotypic analysis are listed in the inset (right). **B,** Hit compounds identified in the phenotypic screen. Vorinostat is labeled as SAHA in **A** and **B. C,** SCC FAK-WT, FAK-G431A,F433A, and FAK−/− cells treated with 10 μM vorinostat. Blue, nuclei; green, F-actin; red, HCS Cell Mask. Scale bar, 50 μm. **D,** Combination of vorinostat or panobinostat with VS-4718 inhibits spheroid growth. **E,** Dose matrices of spheroid area following treatment with vorinostat (top) or panobinostat (bottom) in combination with VS-4178 for 7 days. **F,** Quantification of GFP (G2/M phase) and RFP (G1 phase) FUCCI expression in SCC FAK-WT spheroids treated with HDAC inhibitors for 7 days. For combination treatments with VS-4718, 500 nM VS-4718 was used. RFI, relative fluorescent intensity. **G,** SCC FAK-WT spheroids expressing FUCCI cell cycle reporter were treated for 7 days with vorinostat (Vorino; 1.25 μM) or panobinostat (Pano; 6.25 nM) in combination with VS-4718 (500 nM). For **D, E,** and **F,** data are normalized to DMSO values and are displayed as means ± SEM (*n* = 3 independent experiments). *, *P* < 0.05; **, *P* < 0.01; ***, *P* < 0.001 (two-way ANOVA). **H,** Combination of panobinostat and VS-4718 inhibits SCC FAK-WT tumor growth. Mice were treated with drug(s) from day 0. Group tumor volumes are displayed as mean ± SEM [*n* = 5 mice per group (2 tumors per mouse)]. ***, *P* < 0.001 (one-way ANOVA).

Strikingly, the majority of compounds (69.2%; 9 compounds) whose phenotypes were influenced by FAK kinase activity were HDAC inhibitors (Fig. 2B). HDAC inhibitors induced morphological changes that required FAK kinase activity; representative images, while not conveying the detailed readouts of the multiparametric phenotypic analysis performed, do nonetheless display visual phenotypic differences following treatment with the HDAC inhibitor SAHA (trade name, vorinostat (name used from here on)) that is shown as an exemplar (Fig. 2C). In response to treatment with the HDAC inhibitor, FAK-WT-expressing SCC cells developed stress fiber-like F-actin structures and an enlarged cell area compared to untreated cells, which were not evident with the loss of either FAK or its kinase activity alone. Indeed, HDAC inhibitor treatment appeared to induce a distinct spindle cell morphology characterized by long thin cell protrusions in FAK−/− and FAK-G431A,F433A-expressing SCC cells relative to FAK-WT-expressing SCC cells (Fig. 2C). These data highlight the dependency of HDAC inhibition on FAK signaling to induce robust, and visible, phenotypic effects.

We next tested whether the phenotype observed following treatment with the HDAC inhibitor could be replicated in SCC cells expressing FAK-WT by co-treatment with small-molecule inhibitors of FAK and HDAC. We therefore selected vorinostat, the first HDAC inhibitor approved for clinical use, and panobinostat, a very potent HDAC inhibitor currently undergoing clinical trials (reviewed in (19)). Both vorinostat and panobinostat recapitulated the phenotypic response we observed using the FAK-G431A,F433A mutant (Supplementary Fig. S2A) when they were combined with the FAK inhibitor VS-4718, although this was not linked to a significant difference in cell proliferation in 2D cell culture when HDAC inhibitors were used either to treat cells expressing the FAK-G431A,F433A mutant for 24 hours (Supplementary Fig. S2B). SCC cells expressing FAK-WT in the presence of the VS-4718 FAK inhibitor and treated with either HDAC inhibitor resulted in a weakly anti-proliferative effect after 72 hours treatment (Supplementary Fig. S2C) and in keeping with this, did not significantly alter the G1 cell cycle arrest induced by HDAC inhibitors in 2D culture (Supplementary Fig. 2D). However, since we previously showed that FAK activity is required for proliferation of SCC cells only in 3D culture, but not in 2D (6), we next tested whether the phenotypic effects observed were associated with an anti-proliferative combination in 3D culture. We found that combining FAK and HDAC inhibitors significantly and synergistically inhibited the growth and viability of spheroids composed of FAK-WT-expressing SCC cells, as quantified by measuring spheroid area and viability using calcein AM staining (Fig. 2D and E; Supplementary Fig. S2E-G) and resulted in a robust G1 cell cycle arrest (Fig. 2F and G). We assessed the effect of combining FAK and HDAC inhibitors on SCC tumors grown in CD-1 nude mice in vivo. We injected mice on both flanks with 2.5 × 10^5^ SCC cells expressing FAK-WT and treated established tumors from day 0 with VS-4718 (75 mg/kg twice daily by oral gavage) and panobinostat (5 mg/kg, 5 out of 7 days by intraperitoneal injection). We found that mice bearing SCC tumors expressing FAK-WT that were treated with a combination of VS-4718 and panobinostat exhibited significantly reduced tumor growth compared with either drug alone, each of which had a modest effect on tumor growth when used as a single agent (Fig. 2H).

### Identification of Cells that are Sensitive to FAK and HDAC Inhibitor Combinations

To determine whether other cancer cells (other than the SCC cells we had used for the chemical-genetic phenotypic screen) were sensitive to FAK and HDAC inhibitor combinations, we screened a panel of 33 commonly used human cell lines from a variety of cancer types. Cell lines were treated for 24 hours with FAK and HDAC inhibitor combinations before fixation and quantification of the number of Hoechst-labeled nuclei. We surveyed a number of cell lines for synergistic response to the combinations of FAK and HDAC inhibitors by quantifying the number of nuclei following 24 hours of drug treatment (Supplementary Fig. S3A and B), identifying a number of cell lines (5 out of 33 tested) and two of these sensitive cell lines were examined further; namely A549 (lung adenocarcinoma) and Flo1 (esophageal adenocarcinoma). These exhibited a robust reduction in proliferation in response to the combination of VS-4718 with a HDAC inhibitor even in 2D culture (Fig. 3A). The ability of both A549 and Flo1 cells to grow in a colony formation assay over 7 days was visibly reduced by FAK and HDAC inhibitor combinations (Fig. 3B), and also when the cells were grown as spheroids (Supplementary Fig. S3C), when compared to either drug alone. These data confirmed that A549 and Flo1 cells were sensitive to the FAK and HDAC inhibitor combination, in 2D as well as 3D culture. In the case of these cells, the combination of the FAK inhibitor VS-4718 with either vorinostat or panobinostat synergistically reduced cell viability over a 3-day incubation period (Fig. 3C; Supplementary Fig. S4A–C) and significantly altered the cell cycle distribution, shifting cells into the G2/M phase, as quantified by DNA labeling with Hoechst (Fig. 3D). Apoptosis induced by each HDAC inhibitor in A549 and Flo1 cells was significantly increased by combining with the FAK inhibitor VS-4718 (Fig. 3E), and the growth of A549 tumors in CD-1 nude mice after subcutaneous injection was substantially impaired by combination treatment when compared to either treatment alone (Fig. 3F). This showed that, in a sub-set of cancer cell lines, treating with a combination of FAK and HDAC inhibitors has synergistic and potent effects on tumor growth.

**Figure 3.**
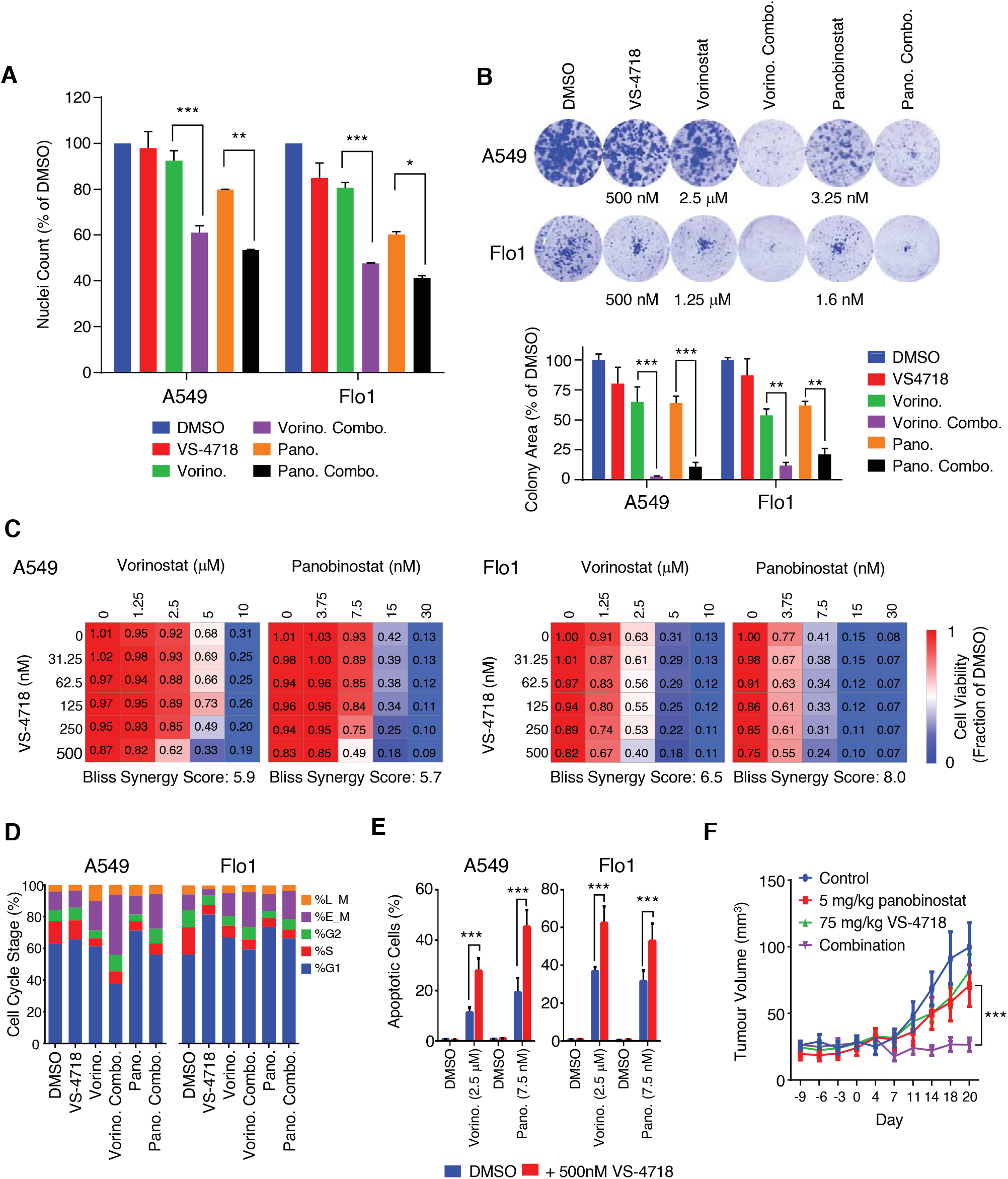
Identification of cells that are sensitive to FAK and HDAC inhibitor combinations. **A,** Nuclei counts of A549 (left) and Flo1 (right) cells following treatment with vorinostat (5 μM), panobinostat (7.5 nM), and VS-4718 (500 nM) for 24 hours. Data are normalized to DMSO values and are displayed as mean ± SEM (*n* = 2 independent experiments). *, *P* < 0.05; **, *P* < 0.01; ***, *P* < 0.001 (two-way ANOVA). **B,** Combined inhibition of FAK and HDAC blocks colony formation. Representative images are shown (top). Bar charts (bottom) are plotted as means ± SEM (*n* ≥ 3 independent experiments). **C,** Combination of FAK and HDAC inhibition reduces cell viability. Dose matrices of viability of A549 (top) and Flo1 (bottom) cells following treatment with vorinostat (μM; left) or panobinostat (nM; right). **D,** Cell cycle distribution in A549 (left) and Flo1 (right) cells following treatment with the same drugs as used in B for 24 hours. Results in C and D are displayed as means (*n* ≥ 3 independent experiments). **E,** FAK and HDAC inhibition enhances the induction of apoptosis in A549 (left) and Flo1 (right) cells. Bar charts are plotted as means ± SEM (*n* ≥ 3 independent experiments). **F,** Combination of panobinostat and VS-4718 inhibits A549 tumor growth. Mice were treated from day 0. Tumor volumes are plotted as means ± SEM [*n* ≥ 4 mice per group (2 tumors per mouse)]. For **B, E,** and **F,** *, *P* < 0.05; **, *P* < 0.01; ***, *P* < 0.001 (one-way ANOVA).

### HDAC and FAK Inhibitors Together Abolish FAK Activation and Promote Nuclear Exclusion of YAP

To understand the signaling events that occurred in cells following treatment with the combination of FAK and HDAC inhibitors, we first assessed FAK-Y397 autophosphorylation. We found that, although the FAK inhibitor VS-4718 was effective at partially inhibiting FAK’s autophosphorylation (and presumed activation – a finding we have made previously in 2D culture (7)) – treatment with both HDAC inhibitors also inhibited FAK-Y397 autophosphorylation to some extent by unknown mechanisms (Fig. 4A). When combined, VS-4718 together with either HDAC inhibitor significantly inhibited FAK-Y397 autophosphorylation to a greater (visibly complete) extent than either VS-4718 or HDAC inhibitor alone.

**Figure 4.**
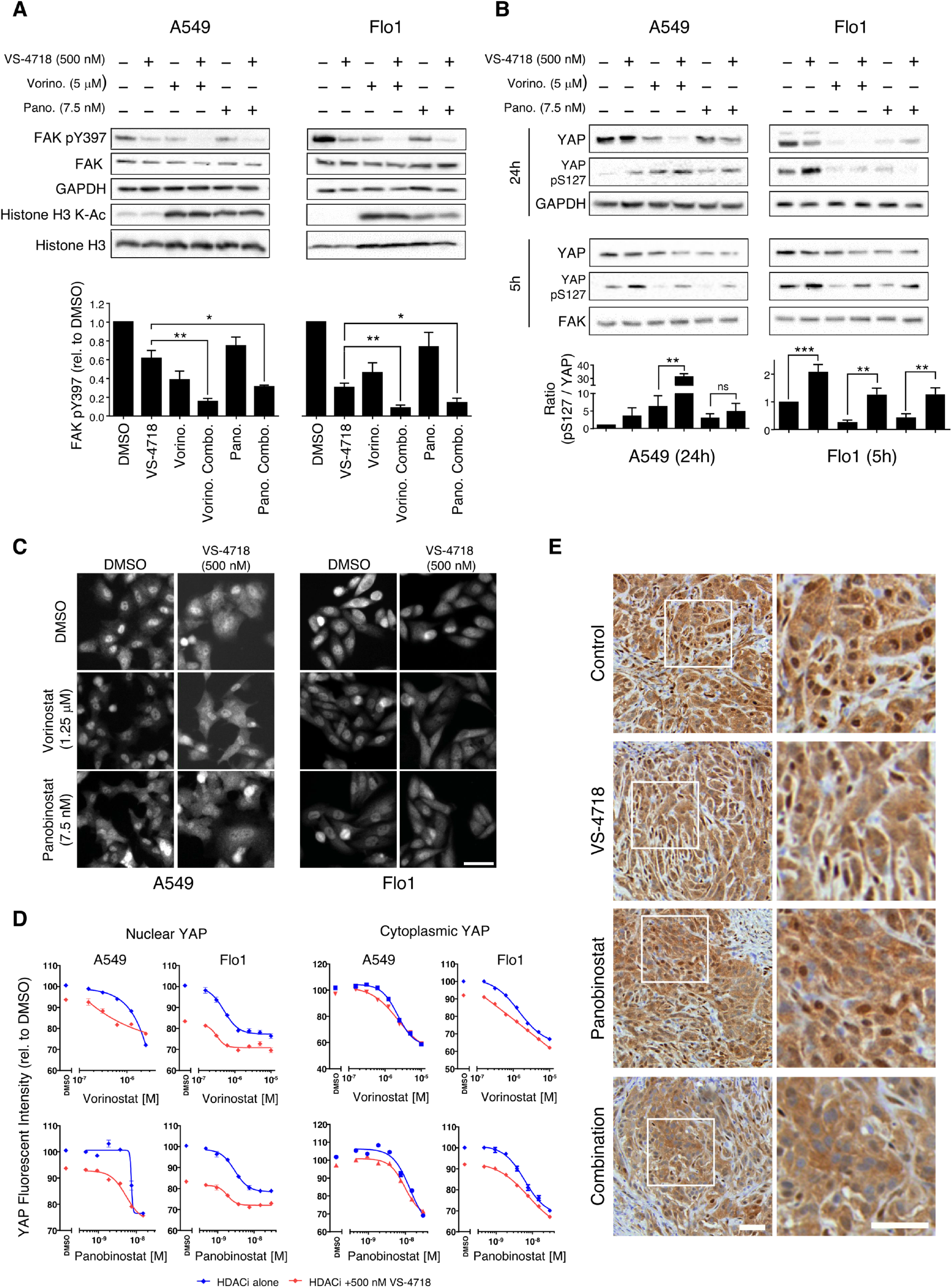
Combined inhibition of HDAC and FAK abolishes FAK kinase activity and regulates YAP localization. **A,** Lysates from A549 (left) and Flo1 (right) cells treated as indicated were immunoblotted for FAK, phosphorylated FAK (Y397), GAPDH, histone H3, and acetylated (K-Ac) histone H3. Quantification of phosphorylated FAK (Y397) immunoblotting is displayed as means ± SEM (*n* = 3 independent experiments). *, *P* < 0.05; **, *P* < 0.01 (one-way ANOVA). **B,** Lysates from A549 (left) and Flo1 (right) cells were immunoblotted for YAP, phosphorylated YAP (S127), and GAPDH at 5 and 24 hours post drug treatment. **C,** A549 and Flo1 cells were fixed and labeled with anti-YAP antibody after 24 hours with drug treatment. Scale bar, 50 μm. **D,** Quantification of nuclear and cytoplasmic anti-YAP labeling. Relative fluorescent intensity values of anti-YAP labeling in the nuclear (left) and cytoplasmic (right) cellular compartments were normalized to DMSO values and displayed as mean ± SEM (*n* = 3 independent experiments). **E,** A549 tumor sections labeled with anti-YAP antibody. Boxed regions are enlarged on the right. Scale bar, 200 μm.

To examine downstream changes in signaling pathways following exposure of Flo1 cells to an effective dose-ratio combination of FAK and HDAC inhibitors relative to each monotherapy, we performed reverse-phase protein array (RPPA) analysis across a number of canonical cancer cell signaling pathways. We found that YES-associated protein (YAP) expression and phosphorylation was altered by the combination (Supplementary Fig. S5). YAP is a co-transcriptional regulator whose activity and nuclear localization is regulated by HDAC activity and FAK signaling (20,21). Therefore, we tested the effect of combined HDAC and FAK inhibitors on YAP. The FAK inhibitor VS-4718 induced YAP phosphorylation on S127, and HDAC inhibitors suppressed YAP protein expression, which was enhanced when the FAK and HDAC inhibitors were combined in both A549 and Flo1 cells (Fig. 4B). Furthermore, in both A549 and Flo1 cells, the FAK inhibitor significantly reduced nuclear localization of YAP, as judged by a reduction in YAP fluorescent intensity in the nucleus (Fig. 4C). Combining FAK and HDAC inhibitors reduced YAP nuclear localization to a greater extent than individual agents alone, as quantified by measurement of fluorescent intensity of anti-YAP cell labeling (Fig. 4D). To determine whether YAP’s localization was sensitive to the combination of FAK and HDAC inhibitors in other cell lines where synergistic anti-proliferative effects between FAK and HDAC inhibitors were not observed, we tested four insensitive cancer cell lines. These did not display the same response with respect to YAP as the sensitive Flo1 or A549 cells, i.e. there was no significant anti-proliferative effects or any visible alteration of YAP’s nuclear localization in response to the combination of FAK and HDAC inhibitors (Supplementary Fig. S7A–C). Two of the non-responsive cell lines, SKGT4 and JH-Eso-AD1, displayed very little nuclear YAP, while the OAC-P4C and MFD1 cell lines displayed robust nuclear YAP localization that was not altered by the combination (Supplementary Fig. S6A). The combination of FAK and HDAC inhibitors in these four cell lines did not enhance anti-proliferative effects (as assessed by nuclear counts) or alter the nuclear localization of YAP, compared to DMSO treatment (Supplementary Fig. S6A–C). Finally, we examined the localization of YAP in vivo using A549 cells, where we observed that YAP’s nuclear localization was reduced in tumors that were treated with the combination of the FAK inhibitor, VS-4718, and the HDAC inhibitor, panobinostat (Fig. 4E). Taken together, these data demonstrate that regulation of YAP correlates strongly with the morphological and synergistic anti-proliferative effects, and acts as a marker of cancer cells sensitive to the HDAC and FAK inhibitor combination.

## Discussion

Here, we describe a completely novel and unbiased approach to identifying useful anti-cancer combinations of small-molecule inhibitors that combines genetic modulation of cancer cells and high-content phenotypic screening to genetically fix one axis of combination screening. Using this screening strategy, we were able to capture phenotypic effects in 2D cell culture models that predicted anti-tumor growth in vivo. We have discovered a novel combination of FAK and HDAC inhibitors that demonstrate synergistic effects in a cancer cell context-dependent manner. This is important because, while FAK is regarded as a high-value druggable target for cancer therapy, it is becoming evident that FAK inhibitors will best be used as part of a clinical combination strategy as FAK is a key component of pathways that promote stress-induced survival signaling, including when cells are treated with putative anti-cancer therapies (9,22). In addition to HDAC inhibitors, which formed the predominant MOA class of compounds that synergized with kinase-defective FAK, we identified a further 18 putative hits representing previously approved drugs that have not yet been pursued (Supplementary Table 2) in addition to hits from a diverse chemical library screen with unknown target classes (not shown). These data imply that there are likely to be a number of useful co-vulnerabilities in cancer cells in which FAK is up-regulated and/or mediates critical survival signaling. Because the FAK and HDAC inhibitor combination was effective in only a sub-set of cancer cells, defined by co-promoted changes in the phosphorylation and nuclear localization of the transcriptional co-activator YAP, it will be important to continue to identify additional collaborating targets and inhibitors which are synergistic with FAK inhibitors.

As regards the value of the multiparametric phenotypic screening approach, we noted that the strongest FAK-dependent cell phenotypes following compound screening in 2D culture of SCC cells were not related to proliferation, but rather changes in morphometric phenotypes. Moreover, these morphometric phenotypes predicted combinations that inhibited tumor growth in mice. More traditional screening strategies generally employ single endpoint analysis, such as cytotoxic screens, where cell viability, or number, are quantified in 2D tissue culture. We have, here, and previously (6), shown that FAK inhibition (either genetic or pharmacological) in SCC cells does not inhibit proliferation or viability in 2D, but does so in 3D and in vivo. In the work presented here, prior knowledge of FAK biology in SCC cells (6) led us to predict that a phenotypic dependency on FAK kinase activity in 2D would prove to be an effective anti-proliferative combination in 3D assays and, potentially, in vivo, thus demonstrating the power of capturing an unbiased and complete description of a compound’s phenotype, including its morphometric properties, to describe its MOA. The wealth of information, and predictive value, contained in the cellular morphological fingerprints following perturbation has been well recognized, and their quantification is increasingly being used as an unbiased method to profile compounds through multiparametric high-content microscopy, coupled to new emerging image-informatics and machine learning solutions (11–13,23–25). Hsp90 and its co-chaperone Cdc37 interact with approximately 60% of the human kinome (16), and thus our novel chemical-genetic phenotypic screening approach represents a broader strategy for rapidly identifying novel kinase inhibitor combinations.

HDACs are a widely studied class of epigenetic regulators that are frequently dysregulated in cancer and so have long been targets of therapeutic research (19). The overall strategy of targeting HDACs for cancer therapy is to revert ‘cancerous’ epigenetic changes towards a more ‘normal’-like cell state and to achieve the reactivation of tumor suppressor pathways, growth arrest, and apoptosis or cell death. These changes undoubtedly will cause cancer cells undue stress, and it may be that FAK is a key component of the pathways that mitigate that stress and permit cancer cell survival. In this regard, we note that cell shape (the regulation of which FAK contributes to via well-recognized effects on actin and adhesion networks) can alter histone acetylation (26) and that gene expression profiling of HDAC inhibitor resistance in human myeloma cell lines is reported to involve actin cytoskeleton-associated genes, and FAK itself, as part of a resistance gene signature (27). This supports the link that we have identified here between the regulation of the cellular epigenetic state by HDAC activity and FAK signaling, revealing a co-dependency in multiple cancer cell lines.

Mechanistically, we have found that the combination of FAK and HDAC inhibitors results in complete abolition of FAK kinase activity, more so than the partial suppression of FAK’s kinase activity that is achieved with a FAK inhibitor alone, suggesting that residual adhesion signaling via FAK is performing a critical survival signal in these cells and that this is linked to growth arrest and apoptosis. Since there is no rational reason why HDAC inhibitors would act directly upon FAK kinase – indeed, shorter periods of incubation with HDAC inhibitors do not affect FAK’s autophosphorylation - we postulate that the effects of HDAC inhibitors are likely indirect, perhaps by regulating expression of gene products that influence FAK’s kinase activity.

It is known that HDAC inhibitors reduce YAP expression (28,29), and FAK, and other focal adhesion proteins, are reported to regulate YAP phosphorylation by LATS1/2 kinase on S127, so preventing its nuclear translocation (20). In our studies, the combination of FAK and HDAC inhibitors collaboratively reinforced the inhibition of YAP activity; firstly, HDAC inhibition reduced YAP expression, and secondly, FAK inhibition impaired the nuclear translocation, and hence the transcriptional activity, of YAP. It is known that the cytoskeleton can directly activate YAP, whereby Rho GTPases become activated by stiff extracellular environments (30), a process in which FAK is implicated (1,21); conversely, YAP expression can upregulate focal adhesion proteins to provide a positive feedback loop (31). Interestingly, FAK is implicated in an integrin α3–FAK signaling axis that drives YAP nuclear localization and proliferation of incisor stem cells (32); this pathway could be active in tumor cells to regulate the proliferation of putative cancer stem cell-like populations, as is reported in triple-negative breast cancer (33). Importantly, FAK and HDAC inhibitors are currently under clinical development across several cancer indications (19,34) and can be taken forward for clinical testing as a putative combination therapy.

## Supporting information

Combined Supp. Figures 1-6

## Acknowledgments

We thank R. Pilkington for assistance with mass spectrometry and K. Macleod for RPPA analysis.

