## Supplementary material for "A Synergistic Anti-Cancer FAK and HDAC Inhibitor Combination Discovered by a Novel Chemical-Genetic High-Content Phenotypic Screen": Combined Supp. Figures 1-6

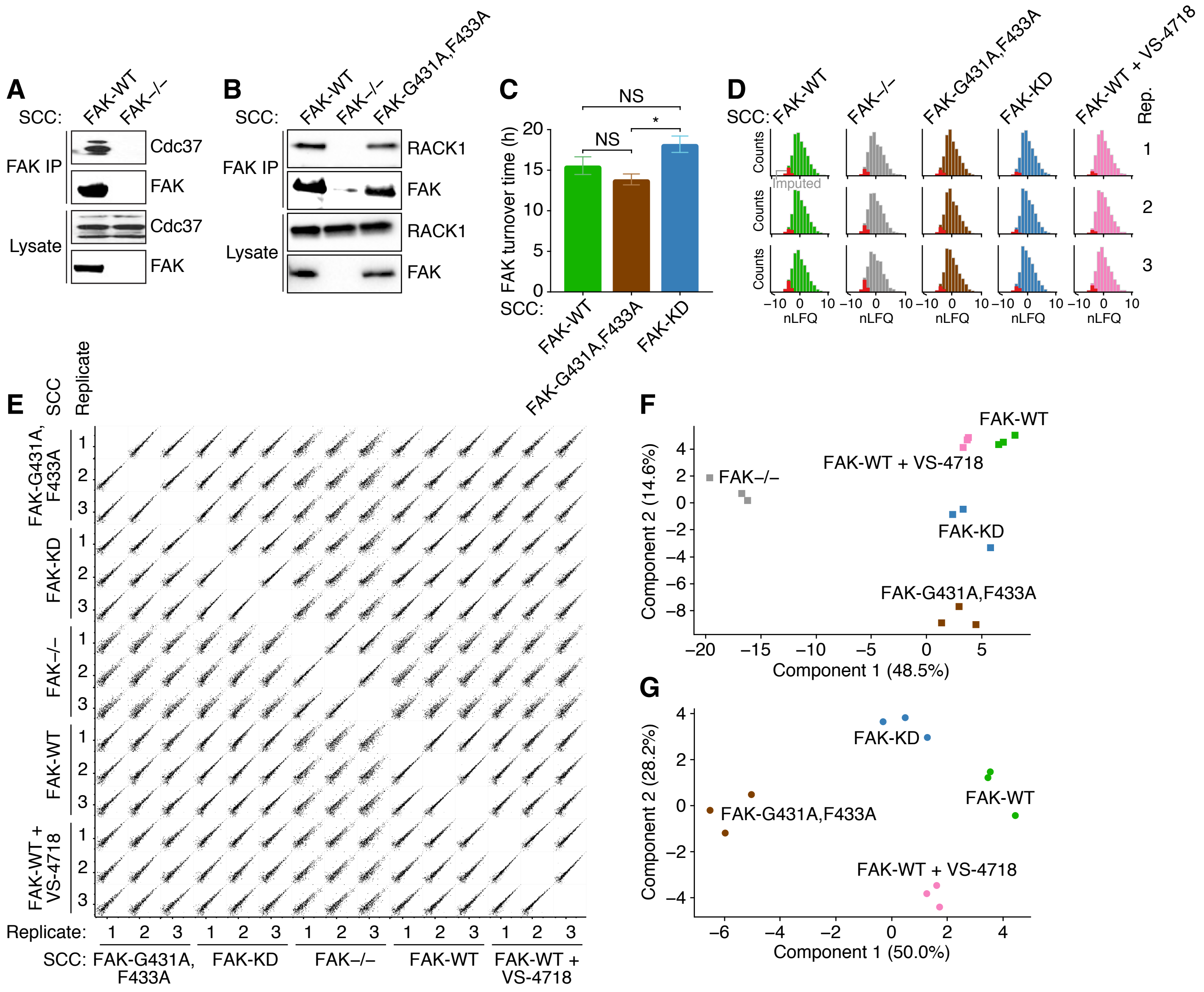


**Supplementary Figure S1.** Proteomic analysis of FAK protein complexes and turnover. **A,** Immunoprecipitation (IP) of FAK from SCC FAK-WT and FAK–/– cell lysates, immunoblotted for FAK and Cdc37. **B,** IP of FAK from SCC FAK-WT, FAK–/–, and FAK-G431A,F433A cell lysates, immunoblotted for FAK and RACK1. **C,** FAK 50% protein turnover times calculated from intersections of normalized SILAC ratio profiles for FAK synthesis and degradation determined by nonlinear regression. No curve best fits in any experiments were statistically significantly different as determined by extra sum-of-squares *F*-test ($\alpha = 0.0083$). Inferred FAK 50% turnover times are plotted as means ± SD ($n = 3$ independent experiments). *, $P < 0.05$; NS, not significant. Statistical significance of turnover times was determined by Kruskal–Wallis test with Dunn's post-hoc test ($\alpha = 0.05$, $H = 6.489$). **D,** Frequency distributions of normalized label-free quantification (nLFQ) values of proteins identified in IPs of FAK from indicated SCC cell lysates by mass spectrometry [false discovery rate (FDR) = 1%]. Missing values imputed from a width-compressed, down-shifted normal distribution are shown in red. **E,** Scatter plots of nLFQ values of proteins identified in IPs of FAK for all pair-wise sample combinations. Significantly differentially regulated proteins as determined by ANOVA (FDR = 5%) are shown in black. **F, G,** Principal component analyses of significantly differentially regulated proteins as determined by ANOVA (FDR = 5%) (**F**) and proteins significantly enriched over control IPs from SCC FAK–/– cell lysates as determined by Student's *t*-test (FDR = 5%) (**G**). The first two principal components account for 63.1% (**F**) and 78.2% (**G**) of the total variance of the respective datasets.

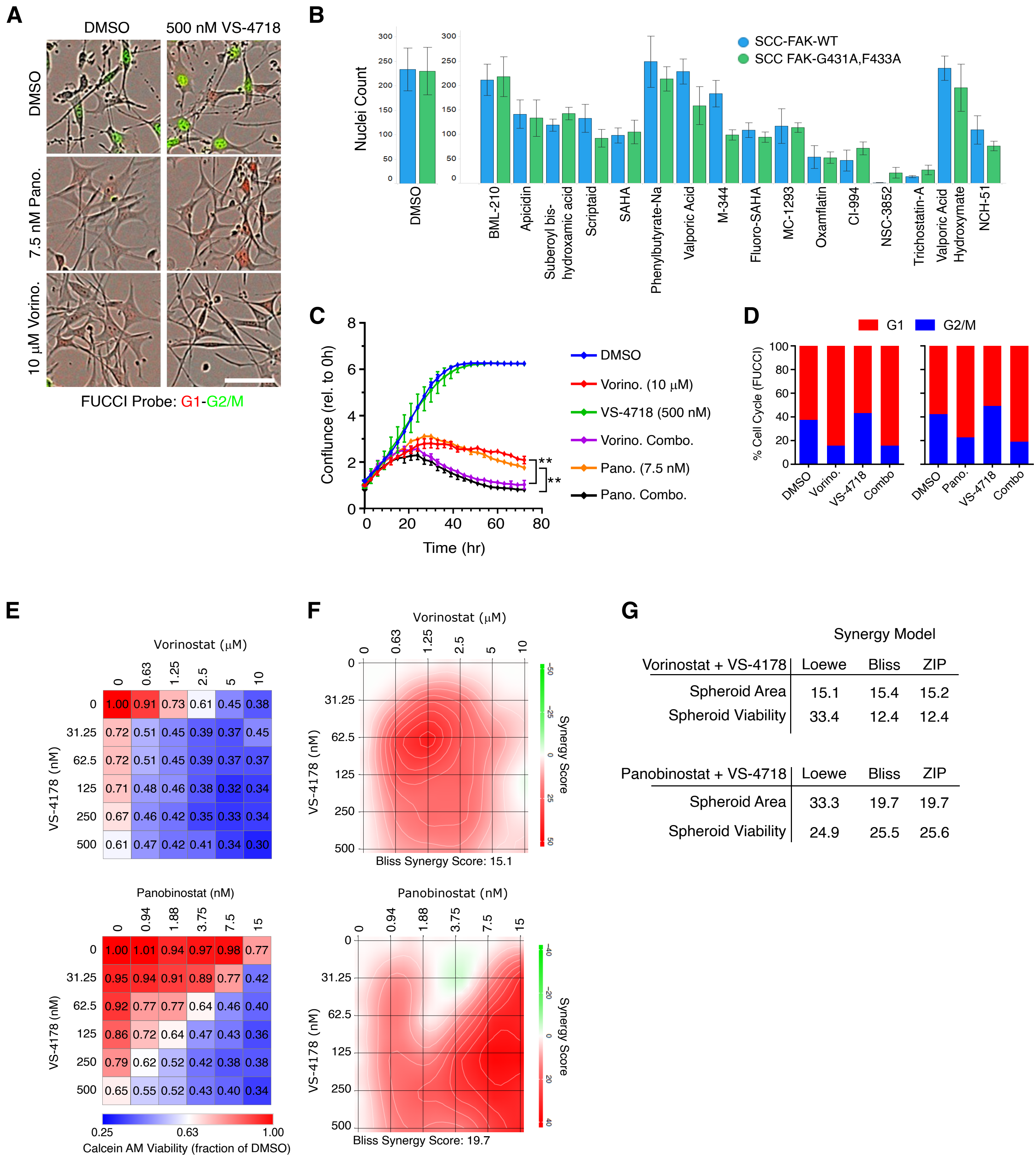


**Supplementary Figure S2.** Analysis of HDAC and FAK inhibition in SCC cells. **A**, SCC FAK-WT FUCCI cells treated with the indicated drugs for 24 hours. Red cells are in the G1 and green are G2/M phases of the cell cycle, respectively. Scale bar is 50 mm. **B**, Quantification of nuclei counts from compound screen. Mean ± SD is shown (n = 6 images). **C**, SCC FAK-WT cell confluence quantified using the Incucyte Zoom microscope. Mean cell confluence is displayed ± SEM (n = 2 independent experiments). NS, not significant. Stastical significance after 72 hours drug treatment was determined by one-way ANOVA followed by a Tukey's multiple comparison test. **, P<0.01. **D**, Quantification of cell cycle stage from FUCCI reporter at 24 hours post drug treatment. Mean population percentages of the cell cycle stage are shown (n = 2 independent experiments). **E**, Drug treated SCC FAK-WT spheroid viability (Calcein AM) following vorinostat (top) or panobinostat (bottom) treatment in combination with VS-4178 for 7 days. (n = 3 independent experiments) **F**, Example Bliss synergy map for SCC FAK-WT spheroid area and treated with vorinostat (top) and panobinostat (bottom) in combination with VS-4718. Calculated from mean spheroid area values shown in Fig. 2E. **G**, Summary table of synergy scores for Bliss, Loewe and ZIP models for HDAC and FAK inhibitor combinations. For **C, D, E** and **F**, data are normalized to DMSO.

A

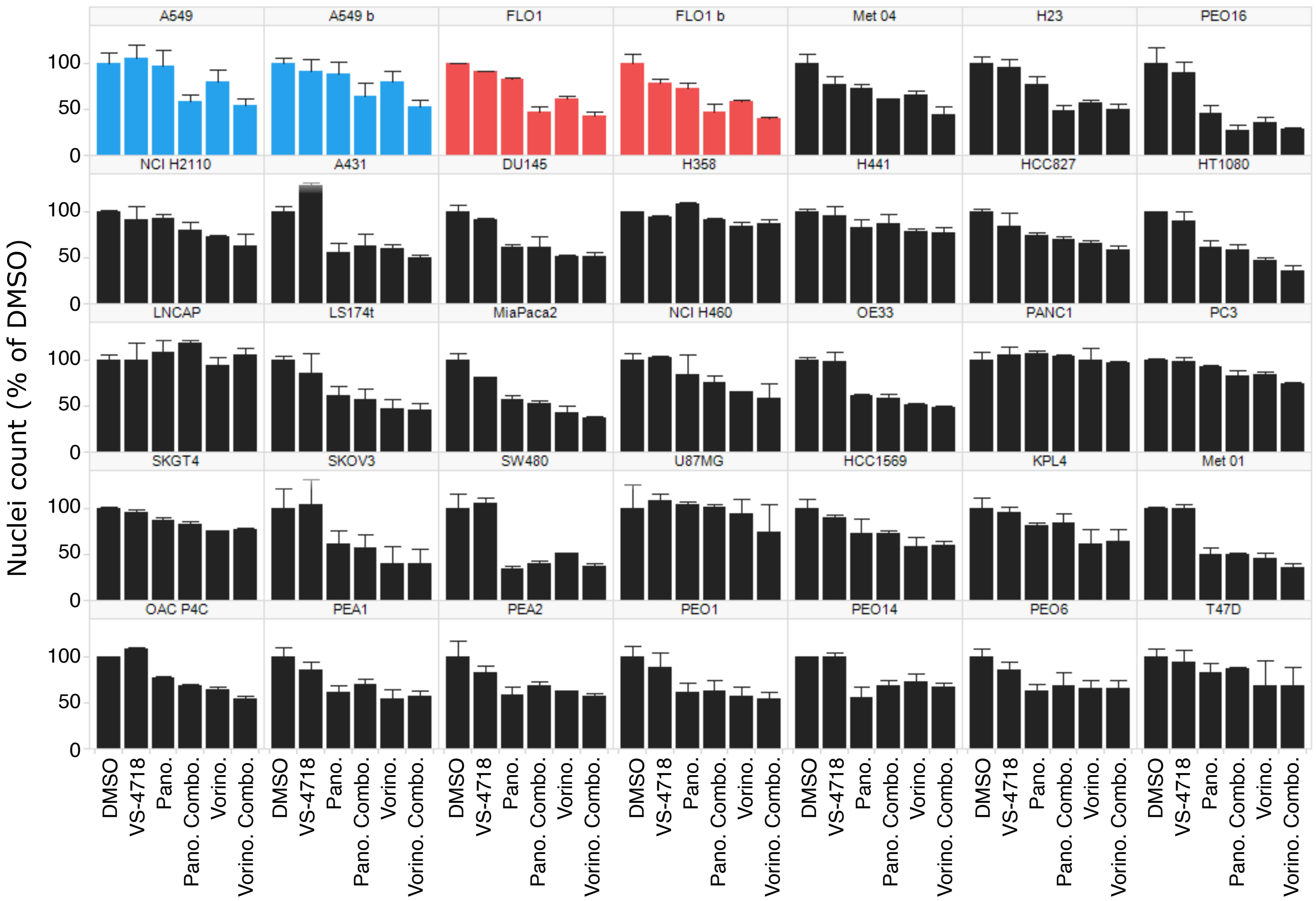


B

| Cell Line | Tumor Type |
|---|---|
| U87MG | Brain |
| HCC827 | Breast |
| T47D | Breast |
| KPL-4 | Breast |
| HCC1569 | Breast |
| Met 01 | Skin |
| Met 04 | Skin |
| A431 | Skin |
| NCI H2110 | Lung |
| NCI H441 | Lung |
| NCI H23 | Lung |
| NCI H358 | Lung |
| A549 | Lung |
| NCI H460 | Lung |
| LS174t | Colorectal |
| SW480 | Colorectal |

| Cell Line | Tumor Type |
|---|---|
| OE33 | Esophageal |
| SKGT4 | Esophageal |
| FLO1 | Esophageal |
| OAC-P4C | Esophageal |
| PEO1 | Ovarian |
| PEO16 | Ovarian |
| PEA1 | Ovarian |
| PEA2 | Ovarian |
| PEO14 | Ovarian |
| PEO6 | Ovarian |
| SKOV3 | Ovarian |
| PANC1 | Pancreas |
| MiaPaca2 | Pancreas |
| PC3 | Prostate |
| LnCap | Prostate |
| DU145 | Prostate |
| HT1080 | Connective |

C

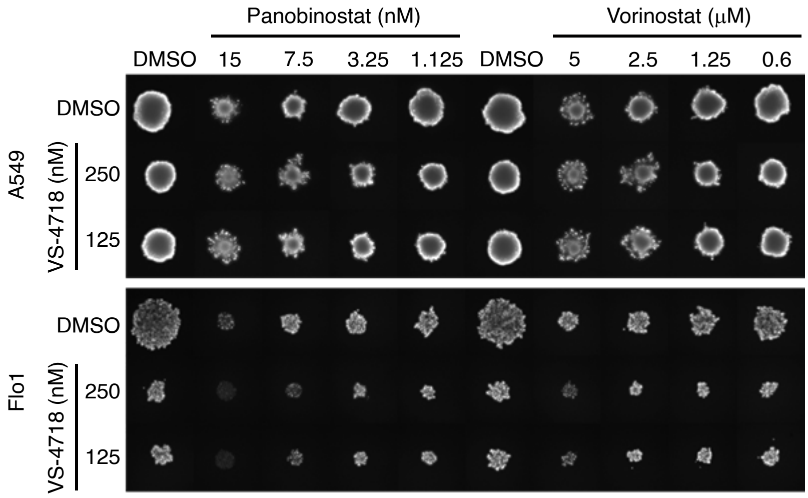


**Supplementary Figure S3.** Analysis of HDAC and FAK inhibition across a panel of 35 human cell lines. **A**, Cells were treated for 24 hours with drugs and Hoechst-labeled nuclei quantified. Drug concentrations used were VS-4718, 500 nM, panobinostat, 7.5 nM and vorinostat, 5 μM. Two replicates for A549 (blue) and Flo1 (red) are shown. Nuclei number are normalized to DMSO and a mean is displayed ± SD (n = 3 replicate wells). **B**, Summary table of human cell lines screened. **C**, A549 and Flo1 cells were cultured as spheroids and treated with drugs for 7 days. Representive images of Calcein AM staining of viable cells in the spheroids are displayed.

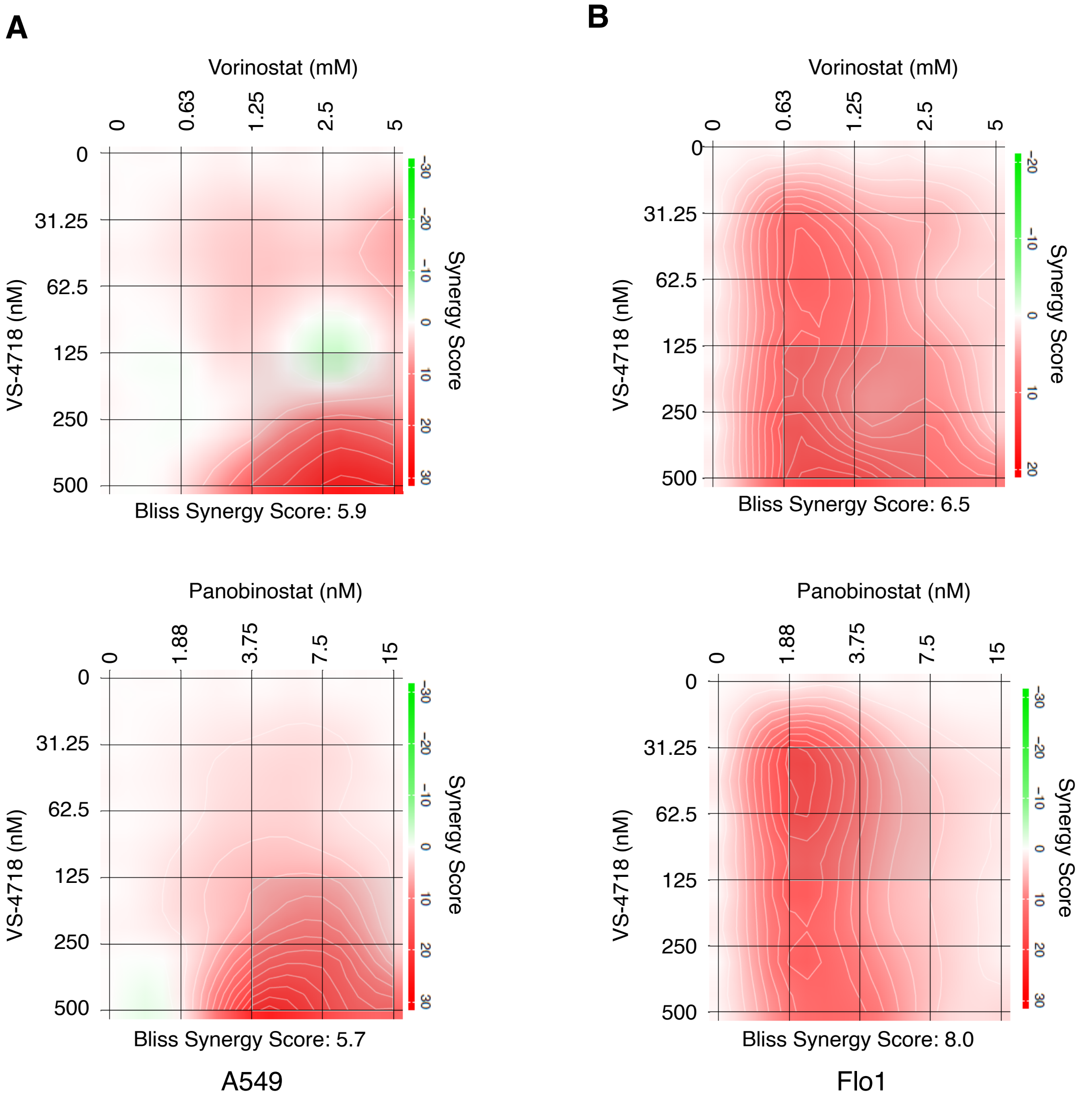


**C**

Synergy Model

| A549 | Bliss | ZIP |
|---|---|---|
| Vorinostat + VS-4718 | 5.9 | 6.1 |
| Panobinostat + VS-4718 | 5.7 | 6.1 |

| Flo1 | Bliss | ZIP |
|---|---|---|
| Vorinostat + VS-4718 | 6.5 | 6.7 |
| Panobinostat + VS-4718 | 8.0 | 8.1 |

**Supplementary Figure S4**. Analysis of HDAC and FAK inhibition in A549 and Flo1 cell lines. **A**, Example Bliss synergy map of A549 cell viability. **B**, Example Bliss synergy map of Flo1 cell viability. For **A** and **B**, data are mean cell viabilities normalized to DMSO from 3-day drug-treated cells as displayed in Fig. 3C, (*n* = 3 independent experiments). **C**, Summary table of synergy models for HDAC and FAK inhibitor combinations.

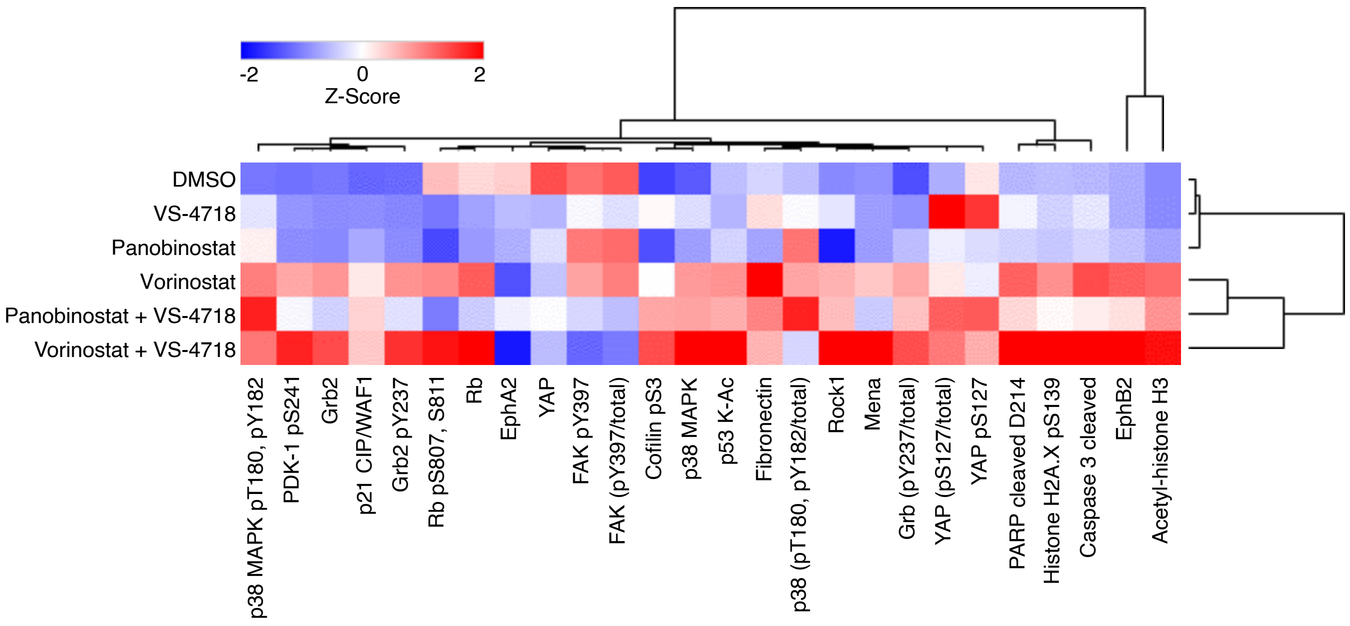


**Supplementary Fig. S5.** Reverse phase protein array (RPPA) pathway analysis. Flo1 lysates were subjected to RPPA analysis using 120 antibodies against canonical cancer cell signaling pathways. Selected antibodies were hierarchically clustered and displayed as a heatmap. VS-4718, 500 nM, vorinostat, 5 μM and panobinostat, 7.5 nM. Mean relative fluorescence value of three technical replicates is shown converted to a *Z*-score.

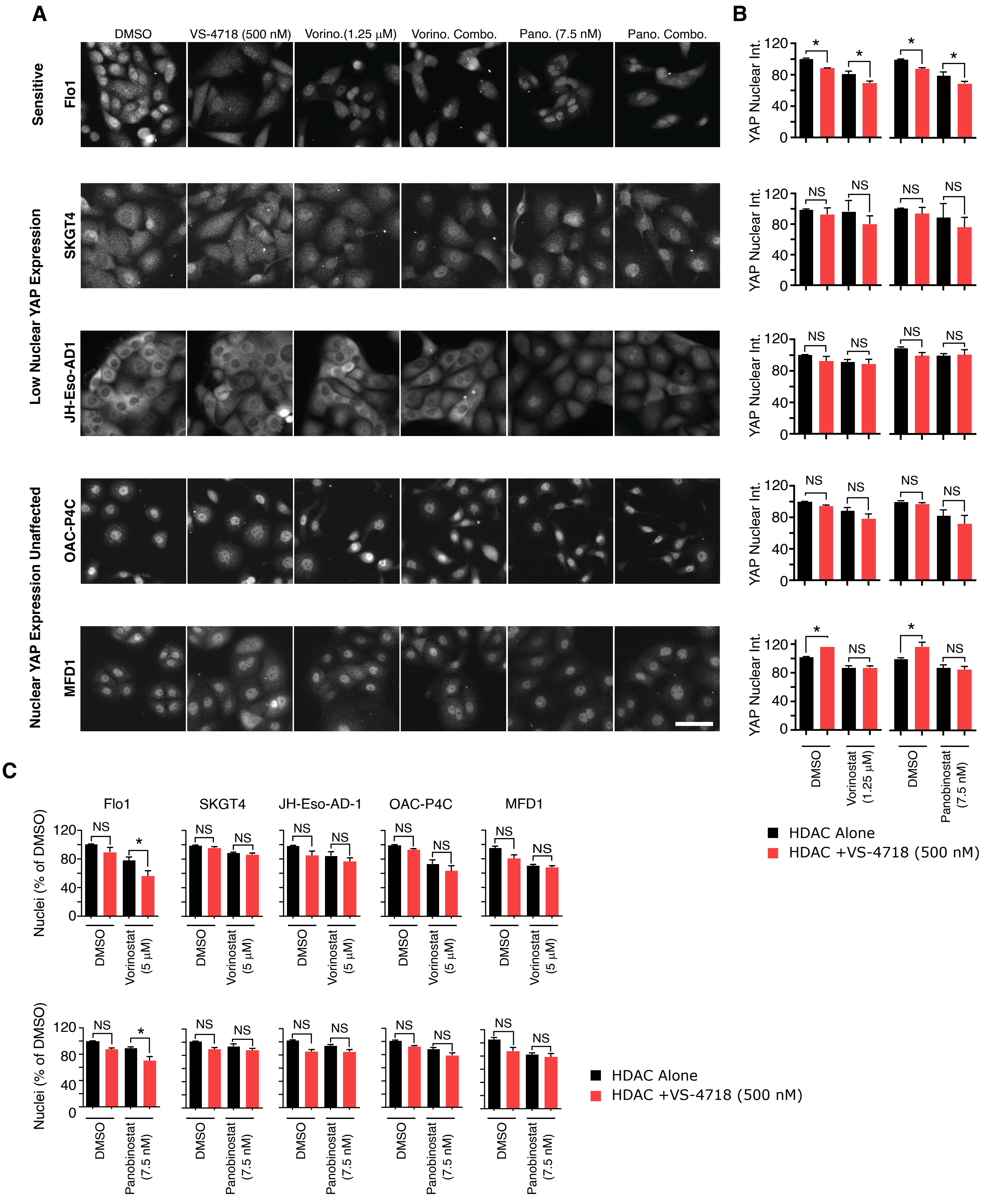


**Supplementary Figure S6.** Quantification of YAP localization in esophageal cell lines. **A**, Immunolocalization of YAP in oesophageal cell lines after 24 hours treatment with indicated compounds. **B**, Quantification of nuclear YAP fluorescence intensity. **C**, Quantification of esophageal cell growth, as measured by nuclei counts, following treatment with compound for 24 hours. Values in **B** and **C** are normalized to DMSO and displayed as mean ± SEM (*n* = 3 independent experiments). Statistical significance was assessed by one-way ANOVA. NS, not significant, *, $P < 0.05$. Scale bar, 50 µm.
